# Paternal inheritance of a vulnerable opioid-taking phenotype in female rats

**DOI:** 10.64898/2026.06.14.732174

**Authors:** Hao Chen, Shuangyin Leng, Sufiya Khanam, Megan K. Mulligan, Eva E. Redei

## Abstract

Risk for opioid use disorder (OUD) is substantially heritable, yet its genetic architecture remains only partly understood. This study examined oxycodone intake in two nearly isogenic rat strains, Wistar Kyoto More Immobile (WMI) and Less Immobile (WLI), and their reciprocal female F1 offspring. The parental strains differ in depression-like behavior and substance use vulnerability, with WMI rats consuming more oxycodone than WLI controls. Voluntary consumption was measured with an operant licking self-administration protocol that delivered 60 μl drug per reward. Across four experimental stages, oxycodone concentrations increased from 0.025 to 0.1 mg/ml, and session durations increased from 1 to 4 hours. Female offspring showed a parent-of-origin effect. F1 females sired by WMI fathers (WLIxWMI) displayed accelerated escalation during the transition from 1-hour to 4-hour sessions in Stage 2 and consumed more oxycodone than reciprocal WMIxWLI females across expanded-access stages. This vulnerability was associated with increased licking during the drug-unavailable timeout period. In WMI and reciprocal WMIxWLI female, consumption was regulated by the drug’s subjective value, as measured by lick microstructure, during Stages 1 and 2. This relationship was absent in WLIxWMI females during Stage 2. Together, these findings suggest that paternal WMI lineage is associated with a rapid transition to high oxycodone intake and cue-directed drug seeking, and identify a parent-of-origin effect that may contribute to female vulnerability to addiction.

## Introduction

Identifying the genetic factors that contribute to Opioid Use Disorder (OUD) is important for addressing the global opioid crisis. Despite evidence of substantial genetic influence (Tsuang *et al*. 1998; Kendler *et al*. 2003), the mechanisms through which parental traits predispose offspring to addiction are often complex and non-linear. Parent-of-origin effects (POE) occur when the behavioral impact of a genetic variant is determined by maternal or paternal transmission. These effects are often driven by genomic imprinting, which can alter neurodevelopmental outcomes and produce sexually dimorphic phenotypes. Investigating these non-canonical inheritance patterns is necessary to resolve how specific genetic backgrounds interact with parental lineage to shape addiction vulnerability.

Reciprocal F1 designs are particularly useful for detecting POE because offspring inherit the same autosomal alleles but differ in the parental origin of those alleles, as well as in associated maternal lineage factors. Prior work in rodents has shown that reciprocal hybrids can differ in anxiety-like behavior, locomotor activity, stress responsivity, stress coping, and drug self-administration, indicating that parental lineage can shape behavior even in the absence of preconception drug exposure (Ahmadiyeh *et al*. 2005; Roy *et al*. 2007; Mont *et al*. 2018; Oreper *et al*. 2018; Chou *et al*. 2022)

This study leverages the phenotypic divergence between two nearly isogenic inbred strains, the Wistar-Kyoto More Immobile (WMI) and Less Immobile (WLI) rats. Originally developed through bidirectional selective breeding to model endogenous depression, the WMI strain exhibits an “internalizing” phenotype characterized by extreme immobility in the forced swim test, diminished exploration in open field environments, and heightened sensitivity to chronic stress (Will *et al*. 2003; Solberg *et al*. 2004). Unlike other preclinical models of stress-induced depression-like behavior, the WMI strain represents an endogenous vulnerability that mirrors the persistent, treatment-resistant symptoms often observed in clinical populations (Andrus *et al*. 2012). Whole-genome sequencing has confirmed that these strains are nearly isogenic, differing by approximately 4,000 homozygous variants (de Jong *et al*. 2021). Clinically, OUD frequently co-occurs with affective disorders, suggesting shared neurobiological vulnerabilities. Previous work established that WMI rats self-administer more oxycodone than WLI rats (Sharp *et al*. 2021), making these isogenic lines a valuable model for exploring the intersection of affective state and OUD risk.

We focused our investigation on female offspring for several reasons. Clinically, women often progress from first drug use to OUD more rapidly than men, a phenomenon known as “telescoping”(Back *et al*. 2010; Greenfield *et al*. 2010; Barbosa-Leiker *et al*. 2021), and report higher rates of comorbid depression and anxiety (Barbosa-Leiker *et al*. 2021). Furthermore, in our previous characterization of the WMI/WLI model, females exhibited significantly more consistent operant licking and higher oxycodone intake compared to males (Sharp *et al*. 2021). These findings suggest that the female reward system may be particularly sensitive to the genetic and epigenetic modulations inherent in the WMI background.

To isolate these genetic influences, we utilized an oral operant self-administration paradigm. In addition to enabling the measurement of voluntary drug intake, this procedure permitted the analysis of lick microstructure, which provides an assessment of the hedonic impact (“liking”) of the solution (Spector *et al*. 1998; Dwyer 2012). By generating reciprocal F1 hybrids (WLIxWMI and WMIxWLI), we aimed to test the hypothesis that POE, which are typically masked in standard genetic designs, contribute to oxycodone intake in female offspring.

## Methods

### Subjects

The study included a total of 51 female rats: WLI (n=13), WLIxWMI F1 (n=13), WMIxWLI F1 (n=15), and WMI (n=10). All rats were young adults, aged 127.1 ± 4.1 days (mean ± SE) at the start of the experiment. All rats were housed under a 12h:12h reversed light/dark cycle (lights on at 9:00 PM and off at 9:00 AM) with food and water available *ad libitum* in their home cages. All procedures were approved by the Institutional Animal Care and Use Committee.

### Operant Licking for Oral Oxycodone Self-Administration

Self-administration was conducted in standard operant chambers (Med Associates) modified for operant licking as previously described (Sharp *et al*. 2021). Each chamber was equipped with two spouts. Licks on the active spout resulted in the delivery of 60 μl of oxycodone solution paired with a 5s cue (tone and light), followed by a 20s time-out period. Licks on the inactive spout were recorded but had no programmed consequences. No prior water restriction or drug priming was used. The procedure consisted of four stages: 0.025 mg/ml in 1h sessions for 3 days (Stage 1), 0.025 mg/ml in 4h sessions for 2 days (Stage 2), 0.05 mg/ml in 4h sessions for 2 days (Stage 3), and finally 0.1 mg/ml in 4h sessions for 10 days (Stage 4).

### Lick Microstructure Analysis

Lick timestamps were recorded with 10 ms resolution using the Med Associates interface. A cluster was defined as a series of licks with inter-lick intervals (ILIs) of less than 0.5s (Davis & Smith 1992). Two primary measures were calculated: the Mean Cluster Size, representing the number of licks per cluster as a proxy for palatability or hedonic value, and the Mean Within-Cluster ILI, representing the average time between licks within a cluster as a measure of motor vigor or reinforcement strength (Spector *et al*. 1998). Outliers were excluded based on physiological limits, specifically an ILI less than 0.12s or greater than 0.22s, and a cluster size less than 2.5 or greater than 20.

### Statistical Analysis

Data were analyzed using Linear Mixed-effects Models (LMM) in R with the lmerTest package. Fixed effects included Strain, Stage, and their interaction, while Animal ID was included as a random effect. For spout discrimination analysis, Spout (Active vs. Inactive) was included as a fixed effect. Degrees of freedom were estimated using Satterthwaite’s method. Significant effects (p < 0.05) were followed by post-hoc Tukey tests using the emmeans package.

## Results

### Acquisition and Spout Discrimination

Female rats across all strains successfully acquired the operant task, demonstrating an ability to discriminate between the active and inactive spouts (Spout: F_1, 1204_ = 1892.92, p < 0.0001; Figure 1). This preference for the active spout was maintained throughout the experiment, with post-hoc analysis confirming significant discrimination (Active > Inactive) for every strain at every experimental stage (all p < 0.0004).

**Figure 1.**
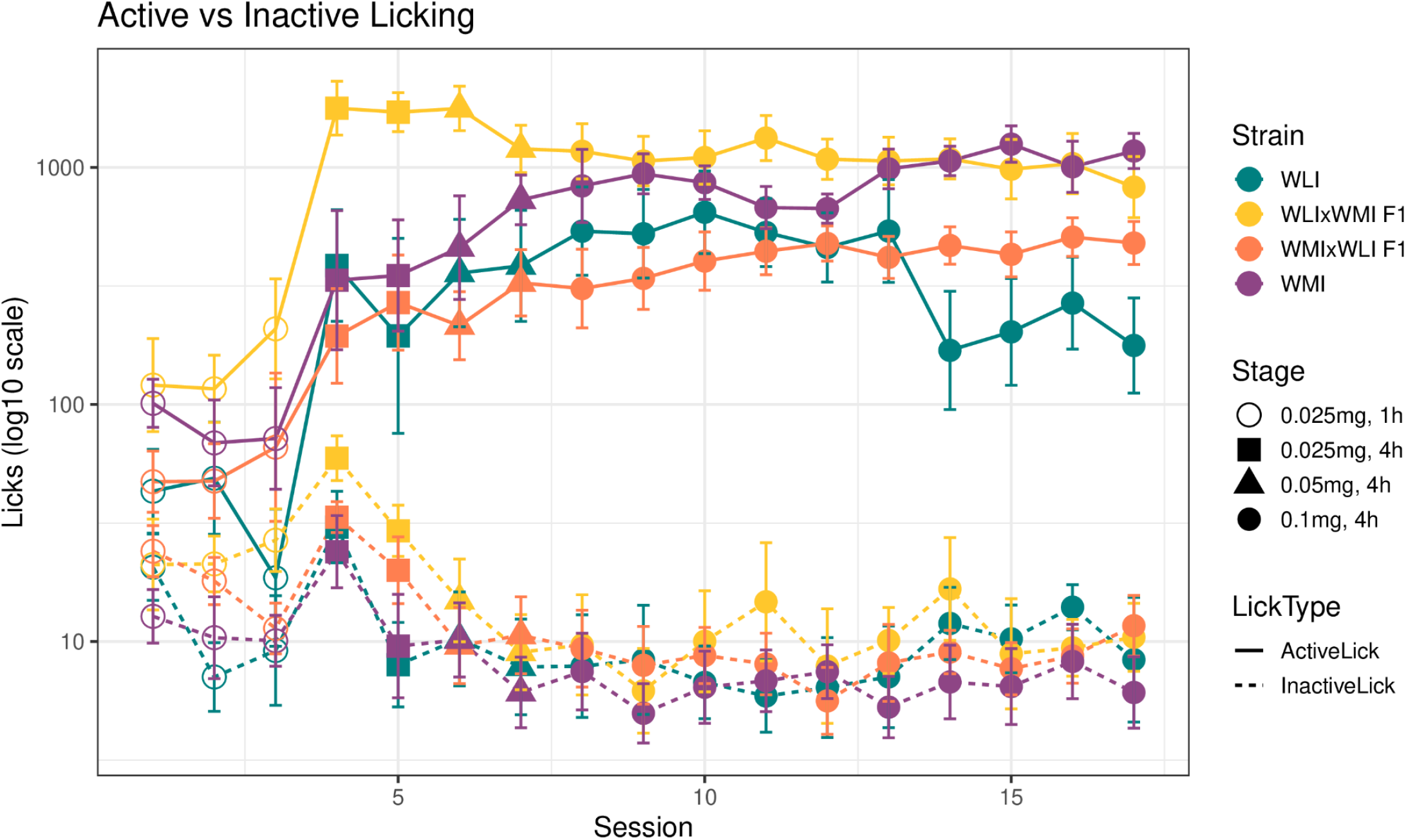
Operant Acquisition and Spout Discrimination. Mean active and inactive licks (± SE) across the four experimental stages. All rat strains successfully discriminated between the active (oxycodone-paired) and inactive (no consequence) spouts, maintaining a consistent preference for the active spout as drug concentration and session duration increased across experimental stages (p < 0.0001).

### Oxycodone Rewards and Intake

Oxycodone self-administration was quantified using two primary measures: rewards, representing the total number of infusions (60 μl each) earned during a session, and intake, representing the total amount of drug consumed normalized to body weight (mg/kg). While rewards reflect the frequency of operant responding, intake provides a physiologically relevant measure of the cumulative drug exposure.

Oxycodone self-administration increased significantly as the concentration and session duration increased (Stage: F_3, 102_ = 50.30, p < 0.0001 for rewards; F_3, 102_ = 106.57, p < 0.0001 for intake; Figure 2). The overall rewards earned and total drug intake across all strains are summarized in Table 1. The two parental strains exhibited distinct drug-taking profiles. The WMI strain, characterized by its “vulnerable” depressive-like phenotype, showed a trend of earning more rewards than the WLI strain at the final 0.1 mg/ml stage (p = 0.08 for rewards), and consumed significantly more oxycodone (p = 0.0002 for intake). By the final stage, WMI females reached an average intake of 2.44 ± 0.28 mg/kg, nearly double that of the WLI females (1.31 ± 0.39 mg/kg).

**Figure 2.**
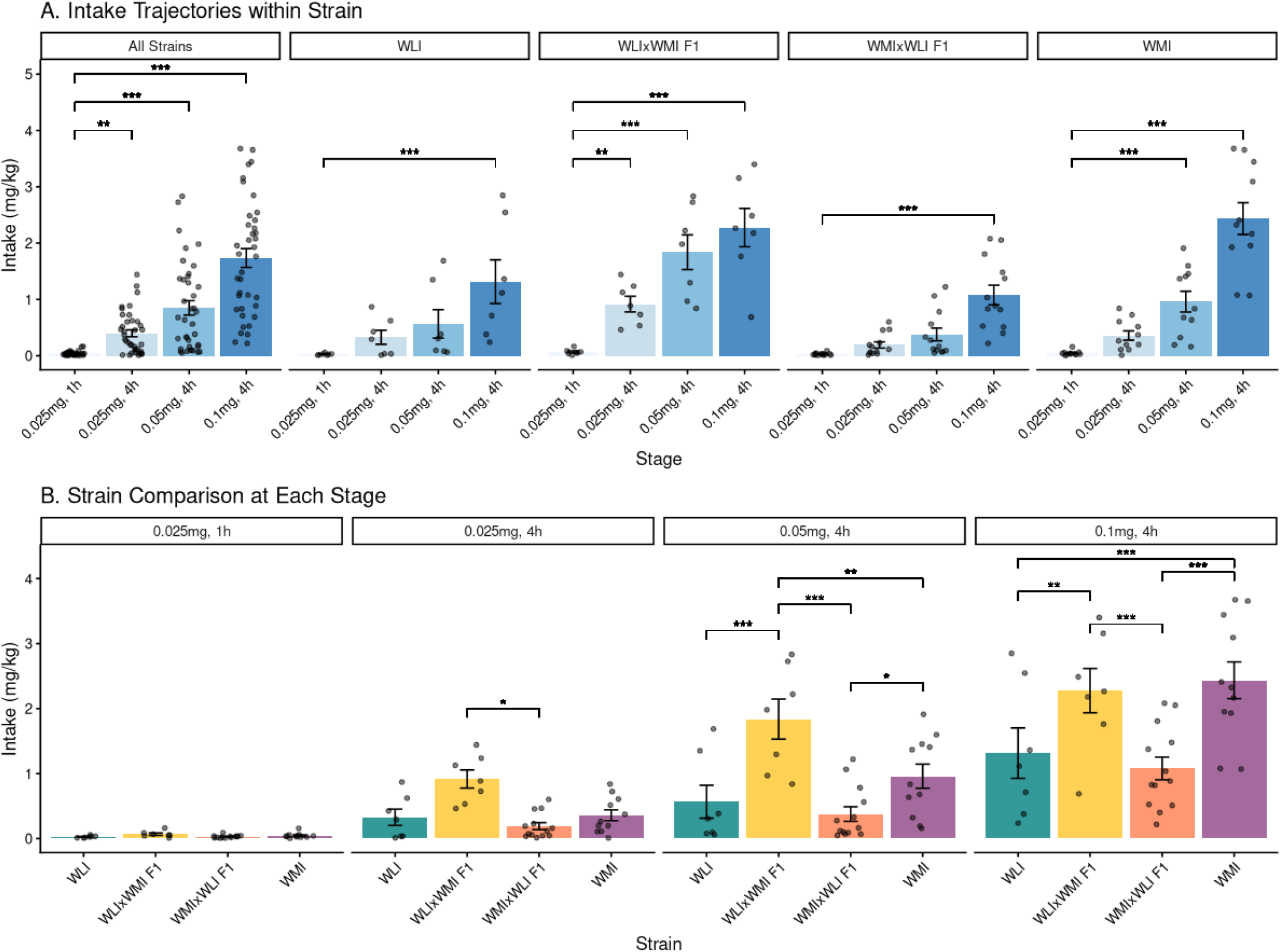
Oxycodone Intake Trajectories and Strain Comparisons. **A)** Intake trajectories (mg/kg, mean ± SE) within each strain across the four experimental stages. WLIxWMI F1 females demonstrated a significant rise in intake as early as Stage 2 (0.025mg, 4h). **B)** Between-strain comparisons of mean intake at each experimental stage. By Stages 3 and 4, WLIxWMI F1 and parental WMI females exhibited significantly higher intake than WLI and WMIxWLI reciprocal hybrids. *p < 0.05, **p < 0.01, ***p < 0.001.

**Table 1.** Summary of Oxycodone Self-Administration by Stage (All Strains Combined)

| Stage | Concentration (mg/ml) | Session Length (h) | Rewards (n) | Intake (mg/kg) |
| --- | --- | --- | --- | --- |
| 1 | 0.025 | 1 | $5.68 \pm 0.82$ | $0.041 \pm 0.006$ |
| 2 | 0.025 | 4 | $47.81 \pm 6.66$ | $0.399 \pm 0.061$ |
| 3 | 0.05 | 4 | $54.09 \pm 7.68$ | $0.850 \pm 0.127$ |
| 4 | 0.1 | 4 | $62.27 \pm 6.04$ | $1.735 \pm 0.166$ |

Data are presented as **Mean ± SE**

A profound POE emerged in the drug-taking behavior, specifically as the drug concentration and session duration increased (Strain x Stage interaction: F_9, 102_ = 6.67, p < 0.0001 for intake). Most notably, WLIxWMI F1 females (paternal WMI) were uniquely identified by an “accelerated intake” phenotype, demonstrating a significant jump in oxycodone intake relative to baseline (i.e., Stage 1) as early as Stage 2 (0.025 mg/ml, 4h) (p = 0.0020; Figure 2). This accelerated trajectory resulted in WLIxWMI females consuming significantly more oxycodone than reciprocal WMIxWLI females during Stage 2 (p = 0.0263), Stage 3 (p < 0.0001), and Stage 4 (p < 0.0001). This early divergence indicates that WLIxWMI females were more susceptible to rapid intake escalation when session constraints were relaxed.

### Transition Sensitivity

To further investigate the increase in intake observed in WLIxWMI F1 females during the first 4-hour access stage (Stage 2), we analyzed “transition sensitivity” to isolate the impact of session duration from drug concentration. By comparing the change in intake during the transition from 1-hour (Stage 1) to 4-hour (Stage 2) sessions, both at the 0.025 mg/ml dose, we quantified the behavioral response to increased drug access. WLIxWMI F1 females exhibited a significantly greater increase in consumption compared to both reciprocal WMIxWLI hybrids (p < 0.0001) and the paternal WMI strain (p = 0.001; Figure 3). While the parental WLI and reciprocal WMIxWLI females showed minimal change in intake relative to baseline (+0.09 mg/kg), WLIxWMI females demonstrated a substantial increase (+0.78 mg/kg). These results suggest that the paternal WMI lineage is associated with sensitivity to the environmental opportunity for increased intake provided by long-access sessions.

**Figure 3.**
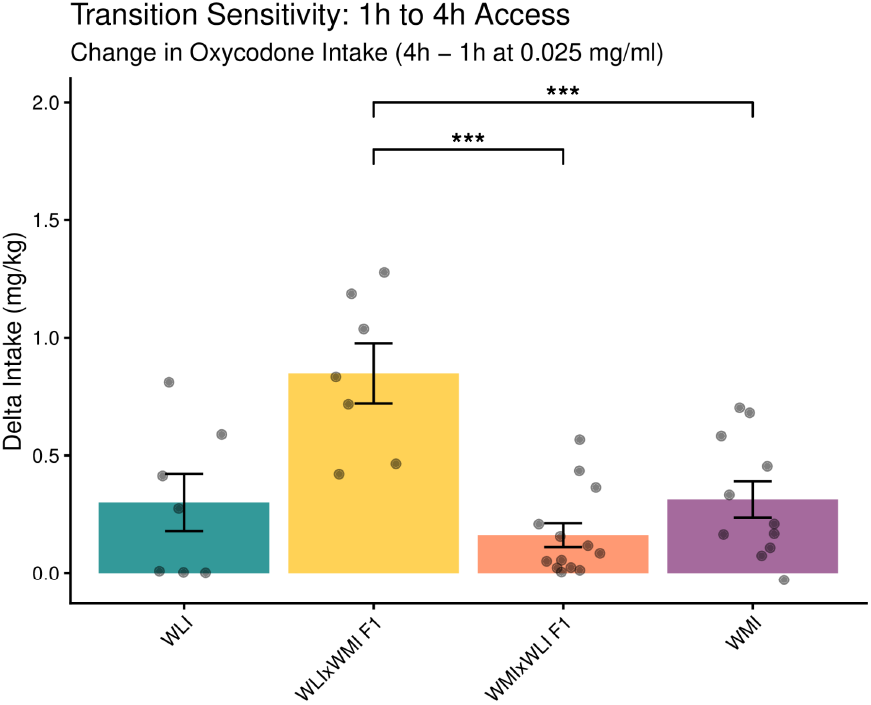
Transition Sensitivity to Session Duration. Mean change in oxycodone intake (mg/kg, ± SE) during the transition from 1-hour (Stage 1) to 4-hour (Stage 2) access sessions at the 0.025 mg/ml dose. WLIxWMI F1 females exhibited a substantially greater increase in intake compared to the reciprocal hybrids and parental WMI strains (p < 0.001), indicating a specific hypersensitivity to the environmental opportunity for increased access.

### Timeout Licking

The increased intake observed in WLIxWMI F1 females prompted an investigation into whether this behavior was driven by deficits in inhibitory control or heightened incentive salience. To assess this, we analyzed the number of active licks during the 20-second post-reward timeout period, a phase when drug delivery is unavailable (Figure 4). Analysis of total timeout licks on the active spout revealed a significant POE, with WLIxWMI F1 females exhibiting a “perseverative” phenotype (Strain: F_3, 34_ = 4.19, p = 0.0125; Stage: F_2, 68_ = 5.52, p = 0.006). During the initial 4-hour stage (0.025 mg/ml), WLIxWMI females performed significantly more non-reinforced licks than their reciprocal WMIxWLI counterparts (1568 ± 387 vs. 463 ± 214 licks, p = 0.0011; Figure 4). This significantly elevated level of timeout activity in WLIxWMI hybrids persisted through the 0.05 mg/ml stage compared to the reciprocal group (p = 0.0142). Notably, the number of licks on the inactive spout during the timeout period remained extremely low (Figure 4B). This contrast between active and inactive spout activity is consistent with an increase in the incentive salience of drug-paired cues, rather than impaired inhibitory control, is a primary contributor to the enhanced oxycodone consumption associated with the paternal WMI lineage.

**Figure 4.**
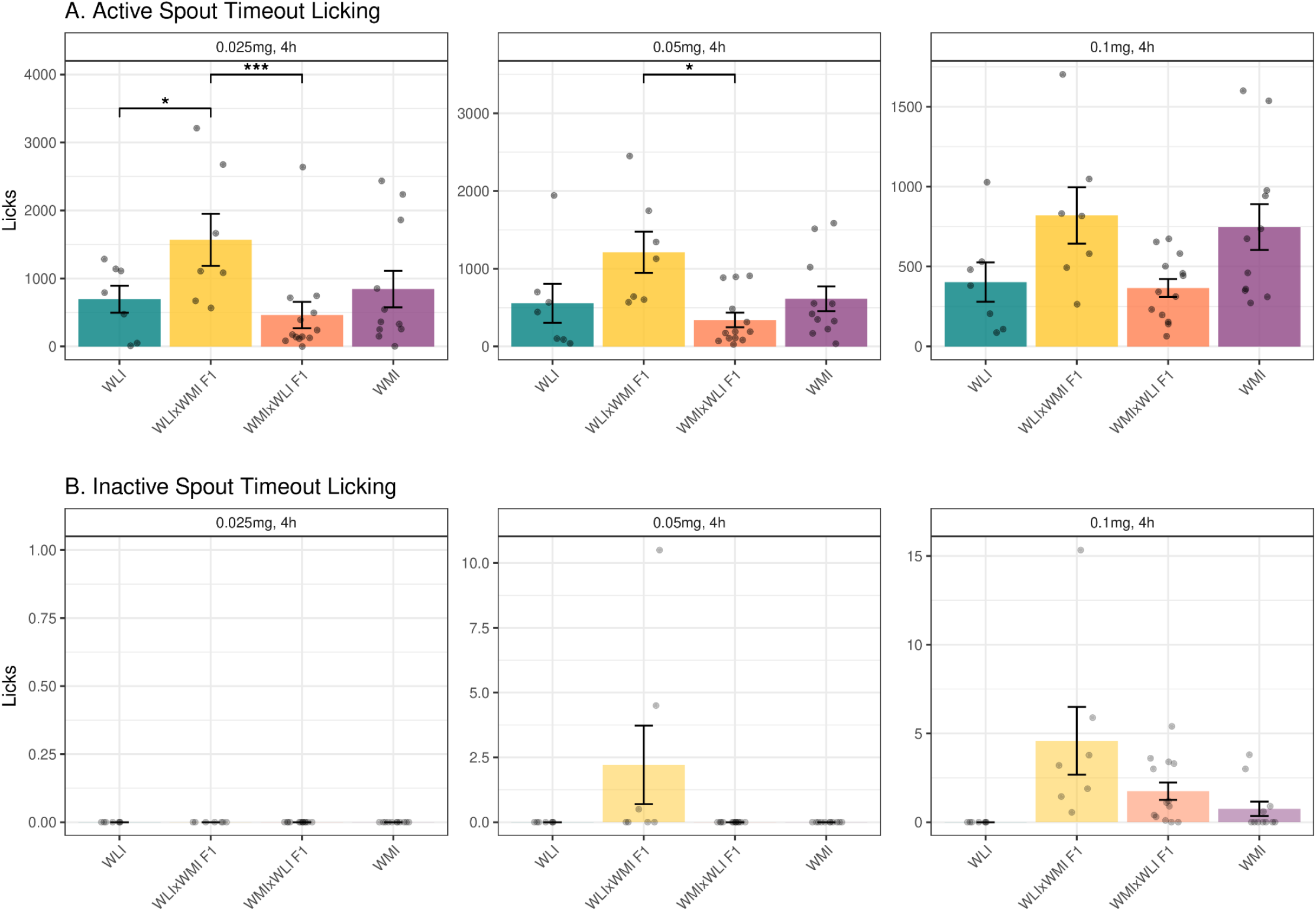
Non-reinforced Licking During Timeout. Total number of licks (mean ± SE) performed during the 20-second post-infusion timeout period on the active (A) and inactive (B) spouts. WLIxWMI F1 females performed significantly more non-reinforced licks than reciprocal WMIxWLI hybrids on the active spout during the first two 4-hour access stages, indicating a surge in incentive salience.

### Lick Microstructure

Lick microstructure analysis was employed to evaluate the hedonic impact (“liking”) across the experimental stages (Figure 5). Analysis of licking patterns revealed dynamic changes across the experiment. When collapsing across strains, there was a trend toward larger clusters as the dose increased (F_3, 111_ = 2, 58, p = 0.0569; Figure 5A), with cluster size showing a Stage by Strain interaction (F_9, 102_ = 2.65, p = 0.0084; Figure 5C). At the initial 4h stage (Stage 2: 0.025 mg/ml, 4h), the parental WMI strain exhibited significantly larger clusters than the parental WLI strain (p = 0.0368; Figure 5C).

**Figure 5.**
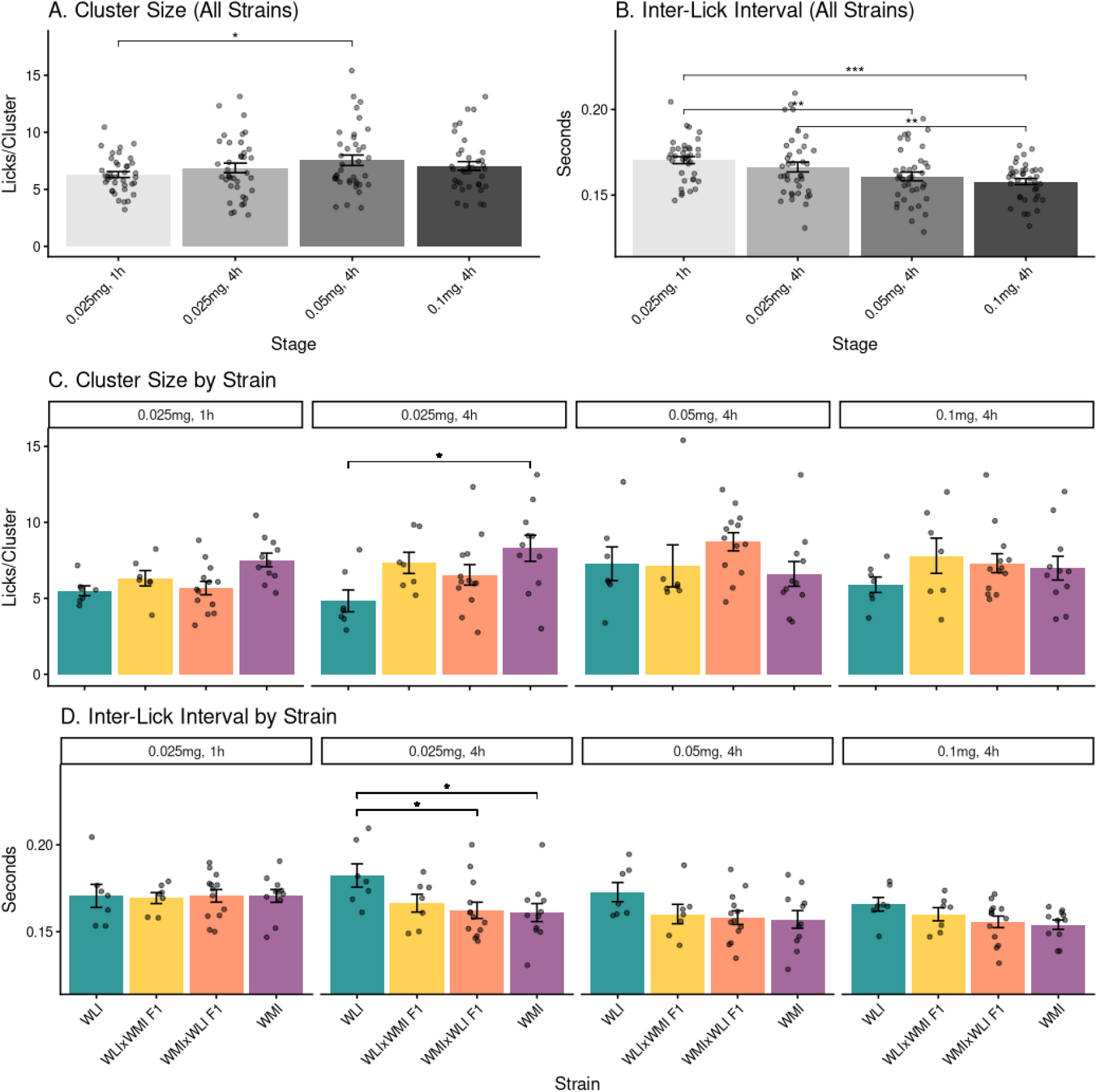
Lick Microstructure: Trajectories and Strain Comparisons. **A-B)** Trajectories for cluster size and inter-lick intervals (ILI) across experimental stages collapsed across all strains. **C-D)** Between-strain comparisons for cluster size and ILI at each experimental stage. Cluster size increased with dose, while ILI decreased. Parental WMI rats exhibited higher cluster size and smaller ILI at the transition to long-access (Stage 2) than WLI rats. Mean ± SE.

The time between licks within a cluster (i.e., ILI) also varied significantly throughout the experiment, primarily decreasing as the concentration of oxycodone increased (Stage: F_3, 111_ = 8.92, p < 0.0001; Figure 5B). This acceleration of lick rate at higher doses was confirmed by post-hoc comparisons, which showed that the ILI at Stage 1 (0.170s) was significantly longer than at Stage 3 (0.161s, p = 0.0027) and Stage 4 (0.158s, p < 0.0001). At the transition to 4 h sessions (Stage 2), the parental WLI strain exhibited a significantly longer ILI (slower licking) compared to both the WMI (p = 0.0016) and WMIxWLI F1 (p = 0.0022) groups, suggesting a lower initial reinforcement strength in the WLI background (Figure 5D). Despite these stage-specific effects, there was no significant main effect of Strain on ILI across the entire experiment (F_3, 34_ = 2.53, p = 0.0736).

To further investigate whether the drug’s hedonic impact contributes to total intake at a finer scale, we performed Pearson correlations between mean cluster size and oxycodone intake for each strain. A critical divergence in behavioral structure was identified during the initial 4-hour transition phase (Stage 2: 0.025 mg/ml; Figure 6). While the parental WMI (r = 0.55, p = 0.012) and reciprocal WMIxWLI F1 hybrids (r = 0.66, p = 0.0004) maintained strong positive correlations between liking and taking, this relationship was not significant in the high-intake WLIxWMI F1 females (r = 0.34, p = 0.24) or the parental WLI strain (r = -0.01, p = 0.98). This indicates that by Stage 2, the increased intake in WLIxWMI hybrids has already become independent of the drug’s immediate palatability. This early-phase decoupling suggests a shift where the hedonic experience no longer serves as a regulatory feedback signal to limit intake, but instead may function as a feed-forward trigger for further consumption. At higher drug concentrations (0.05 and 0.1 mg/ml), these correlations became non-significant across all groups (all p > 0.14), likely due to the potent reinforcing properties of the drug overriding individual differences in hedonic feedback.

**Figure 6.**
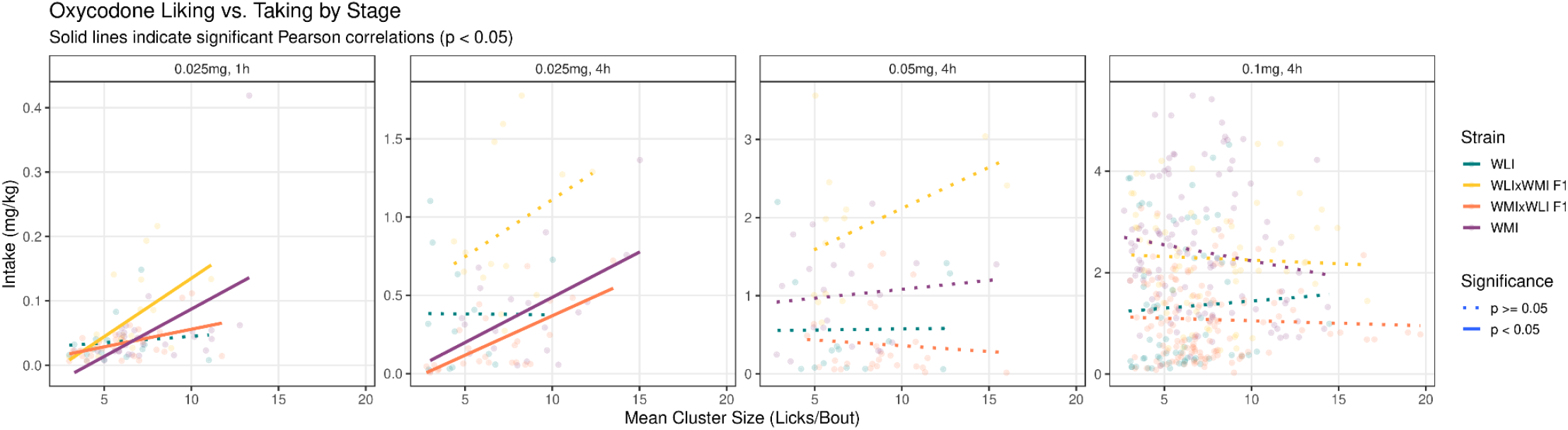
Decoupling of Reward Valuation and Consumption. Pearson correlations between mean cluster size (“liking”) and total oxycodone intake at Stage 2 (0.025 mg/ml, 4h). While intake remained coupled to reward valuation in WMI and reciprocal hybrids (p < 0.05), this relationship was absent in WLI and WLIxWMI F1 females, suggesting an early-phase shift toward palatability-independent drug seeking.

## Discussion

This study identified a POE on the behavioral patterns of oxycodone self-administration that emerged during the initial phases of drug exposure. F1 females inheriting WMI alleles from the paternal line (WLIxWMI) exhibited phenotypes that were latent during the 1 hour sessions of Stage 1. However, the phenotype emerged when session duration was expanded to 4 hours (Stage 2), suggesting that the WLIxWMI cross is sensitive to the environmental opportunity for escalated consumption. This behavioral profile was characterized by increased consumption, enhanced drug seeking during timeout period, and an accelerated shift toward drug seeking independent of palatability. These findings indicate that WMI paternal lineage is associated with susceptibility to high intake states and offer a novel opportunity to evaluate the genetic and epigenetic mechanisms underlying this vulnerability, providing a model for the rapid intake escalation observed in populations at risk for opioid use disorder.

The accelerated intake reflects a reorganization of drug seeking behavior. WLIxWMI females performed more non reinforced licks during the timeout period, indicating heightened engagement with the drug paired spout when oxycodone was unavailable. Lick microstructure provides a second, independent readout of this shift. Cluster size is commonly interpreted as an index of palatability or hedonic value because rats organize licking into clusters whose mean size increases with the concentration of palatable sucrose solutions, whereas manipulations of deprivation primarily increase the number of clusters or meal duration rather than cluster size (Davis & Smith 1992; Spector *et al*. 1998; Dwyer 2012). Dwyer (2012) further argues that lick cluster size is useful for assessing hedonic reactions in rodents because it tracks both unconditioned palatability and learned changes in flavor preference or aversion. In the present study, this hedonic signal predicted oxycodone intake in WMI and reciprocal WMIxWLI females during the transition to 4 hour access, suggesting that consumption in these groups remained coupled to immediate reward valuation. By contrast, the cluster size-intake relationship was absent in WLIxWMI females. Because this decoupling occurred despite strong drug seeking and elevated consumption, it suggests a transition to a high intake state in which oxycodone taking is no longer constrained by immediate palatability. This pattern is consistent with evidence that drinking microstructure can reveal drug-induced changes in hedonic evaluation during operant self administration (Robinson & McCool 2015).

The lineage effects observed here extend prior evidence that reciprocal F1 hybrids can show POE-dependent behavioral differences. Such effects are often domain specific rather than global. For example, reciprocal hybrids of Roman High Avoidance and Low Avoidance strains differ in coping behaviors, such as time spent in open sections of a zero maze, but not in shuttle box avoidance (Mont *et al*. 2018). In that model, offspring phenotypes for anxiety related traits resembled the maternal strain (Mont *et al*. 2018). Similarly, reciprocal hybrids of NOD and C57BL6 strains differ in locomotor activity, with NOD maternal hybrids being more active than F1 offspring with C57BL/6 mothers (Oreper *et al*. 2018).

In contrast to the behaviors above, our findings showed that the paternal lineage dictates oxycodone intake in offspring. This paternal dominance is strongly mirrored in nicotine self-administration, where Fischer 344 sires produce offspring with significantly higher intake (Spence *et al*. 2020; Kozlova *et al*. 2021). Similarly, in models of alcohol preference, offspring consumption levels follow the father’s genotype rather than the mother’s, an effect that persists across multiple generations (Spence *et al*. 2020).

Although the present study used drug-naive parental strains, parental exposure models provide convergent evidence that paternal history can influence offspring drug consumption. Paternal morphine exposure enhances opioid vulnerability in offspring compared to saline controls, with female progeny of morphine-treated sires showing increased morphine intake during acquisition and higher motivation for oxycodone (Vassoler *et al*. 2020). Paternal heroin experience also increases heroin taking and seeking in male offspring compared to yoked controls (Gao *et al*. 2025). These exposure models demonstrate that paternal experience can also bias offspring toward escalated drug consumption.

Together, findings from drug-naive reciprocal crosses and parental exposure models suggest a broader principle: paternal lineage can influence offspring vulnerability through inherited regulation of reward circuitry. In naive reciprocal-cross models, this regulation may reflect genetic background, genomic imprinting, or other POE-dependent mechanisms. In parental exposure models, it may additionally involve drug-induced germline epigenetic changes. Both mechanisms may produce an inherited predisposition that determines voluntary intake through genetic and epigenetic factors (Spence *et al*. 2020).

One potential brain region for these lineage dependent effects is the nucleus accumbens. Reciprocal crosses exhibit parent biased allelic expression. For example, reciprocal hybrids show POE dependent allele specific expression for about 300 genes, many involved in neuromodulation and neural plasticity (Lo *et al*. 2018). Paternal lineage effects in the nucleus accumbens core are also linked to allelic imbalance in genes involved in neurogenesis (Kozlova *et al*. 2021). Paternal opioid phenotypes are further associated with specific regulatory signatures in the accumbens, such as the downregulation of miR 19b (Gao *et al*. 2025). The emergence of the POE during the transition to expanded access suggests a latent regulatory bias. Other evidence indicates that parentage effects in the nucleus accumbens can be magnified by drug exposure itself (Lo *et al*. 2018). The expanded access sessions in this study likely unmasked a pre-existing inherited vulnerability in the WLIxWMI cross.

Identifying the mechanism of this lineage dependent vulnerability requires distinguishing between early environment and inherited factors. The reciprocal cross argues against postnatal WMI maternal care, WMI uterine environment, or WMI mitochondrial inheritance as sufficient explanations for the high intake phenotype. WMIxWLI offspring were gestated and raised by WMI mothers but showed low intake. In contrast, high intake occurred in WLIxWMI offspring, which were gestated and raised by WLI mothers. The parental WLI strain further shows that WLI maternal lineage alone is not sufficient, because WLI females also have a WLI uterine environment, WLI mitochondrial background, and WLI postnatal care yet exhibit low consumption. Thus, the data most strongly support a maternal imprinting (paternal effect) as a plausible biological mechanism, and paternal inheritance can enhance drug consumption through parent specific epigenetic marks (Kozlova *et al*. 2021). However, reciprocal designs cannot definitively isolate canonical imprinting from other parental effects. A permissive contribution of WLI maternal lineage factors, including uterine environment or mitochondrial DNA inheritance, could still interact with paternal WMI lineage (Roubertoux *et al*. 2003; Hager *et al*. 2008). These reciprocal differences therefore reflect a regulatory architecture that is inherently sensitive to paternal origin while leaving open whether the mechanism reflects canonical imprinting, paternal germline epigenetic marks, or interaction with maternal lineage factors (Oreper *et al*. 2018).

The expression of this parent-of-origin effect in females raises an important question about sex specificity. Previous work showed that parental WMI females and males consume more oxycodone than WLI rats of the same sex, and that females generally consume more than males (Sharp *et al*. 2021). However, whether reciprocal F1 males show the same pattern is unknown, so we cannot determine whether the paternal WMI effect is female specific or reflects a broader inheritance pattern. If male reciprocal hybrids do not show the same POE, maternally imprinted genes on the X chromosome would become a plausible contributor to the enhanced oxycodone seeking observed in WLIxWMI females. Although the paternal X chromosome is often silenced, several maternally imprinted genes have been identified on the X chromosome (Abdulai-Saiku *et al*. 2025). Testing male reciprocal hybrids will therefore be necessary to distinguish a female-limited imprinting mechanism from a general paternal-lineage effect.

The greater oxycodone intake of WMI females than males and the accelerated intake of the WLIxWMI females provide a preclinical sex-specific model for rapid escalation of opioid use. Women often progress from initial drug use to dependence more rapidly than men (Back *et al*. 2010; Greenfield *et al*. 2010; Barbosa-Leiker *et al*. 2021). In this study, females of WMI paternal lineage, both paternal and F1, showed early transition to high intake when access was expanded, suggesting that inherited parent-sensitive regulation can lower the threshold for dysregulated drug taking. These findings emphasize the importance of non-canonical inheritance patterns in addiction genetics and identify the paternal WMI lineage as a potential entry point for understanding the heritability of opioid use disorder.

